# TMEM120 is a coenzyme A-binding membrane protein with structural similarities to ELOVL fatty acid elongase

**DOI:** 10.1101/2021.06.13.448233

**Authors:** Jing Xue, Yan Han, Hamid Baniasadi, Weizhong Zeng, Jimin Pei, Nick Grishin, Junmei Wang, Benjamin P. Tu, Youxing Jiang

## Abstract

TMEM120A, also named as TACAN, is a novel membrane protein highly conserved in vertebrates and was recently proposed to be a mechanosensitive channel involved in sensing mechanical pain. Here we present the single particle cryo-EM structure of human TMEM120A which forms a tightly packed dimer with extensive interactions mediate by the N-terminal coiled coil domain (CCD), the C-terminal transmembrane domain (TMD), and the re-entrant loop between the two domains. The TMD of each TMEM120A subunit contains six transmembrane helices (TMs) and has no clear structural feature of a channel protein. Instead, the six TMs form an α-barrel with a deep pocket where a coenzyme A (CoA) molecule is bound. Intriguingly, some structural features of TMEM120A resemble those of elongase for very long-chain fatty acid (ELOVL) despite low sequence homology between them, pointing to the possibility that TEME120A may function as an enzyme for fatty acid metabolism, rather than a mechanosensitive channel.

## Introduction

TMEM120A was initially identified as a nuclear envelope transmembrane protein (NET) by proteomics and was originally named as NET29 (Malik et al., 2010; Schirmer et al., 2003). It was suggested to be preferentially expressed in adipose and plays important role in adipocyte differentiation in an earlier study (Batrakou et al., 2015). In a recent follow-up study, the same group demonstrated that adipocyte–specific *Tmem120a* knockout mice causes disruption of fat-specific genome organization and yields a latent lipodystrophy pathology similar to lamin-linked human familial partial lipodystrophy type 2 (FPLD2) (Czapiewski et al., 2021). However, a completely different function has been proposed for TMEM120A in another recent study in which TMEM120A, renamed to TACAN, was shown to be expressed in the plasma membrane of a subset of sensory neurons and function as a mechanosensitive channel involved in sensing mechanical pain (Beaulieu-Laroche et al., 2020). This finding of a potential novel mechanosensitive channel propelled us to pursue the structural and functional studies of human TMEM120A. However, we were unable to reproduce the mechanosensitive activity of TMEM120A expressed in HEK293 or CHO cells, nor do we observe any channel activity in planar lipid bilayer using TMEM120A protein reconstituted in liposomes. Here we present the single particle cryo-EM structure of TMEM120A which exhibits no obvious feature of a channel protein. Instead, TMEM120A shares several common features with the recently determined ELOVL7 structure (Nie et al., 2020), a member of ELOVL family elongases important for the biosynthesis of very long-chain fatty acid. The ELOVL elongases (ELOVL1-7) are ER membrane enzymes that catalyze a condensation reaction between a long-chain acyl-CoA and malonyl-CoA to produce a 3-keto acyl-CoA, free CoA and CO_2_ (Deak et al., 2019; Jakobsson et al., 2006; Leonard et al., 2004; Pereira et al., 2004), which is the first step in the four-step elongation process of very long-chain fatty acid. While we are unable to define the physiological function of TMEM120A in this study, its structural similarity to ELOVL7 leads us to suspect that TMEM120A may function as an enzyme for lipid metabolism rather than an ion channel.

## Results

### 1. Dimeric structure of TMEM120A

Human TMEM120A was expressed in HEK293F cells using the BacMam system, solubilized in LMNG detergent, and finally purified in digitonin detergent as a homo-dimer (Methods). The single particle cryo-EM structure of TMEM120A dimer was determined to the resolution of 3.2 Å (Figure1a-d, Figure 1—figure supplement 1-3 and Figure 1—source data 1). The EM density map is of high quality, allowing for accurate model building for the major part of the protein containing residues 8-69, 80-255 and 261-335 for each subunit. In addition, electron density from a bound ligand is clearly visible within each subunit and can be modeled as a CoA molecule as will be further discussed later (Figure 1e&f).

**Figure 1.**
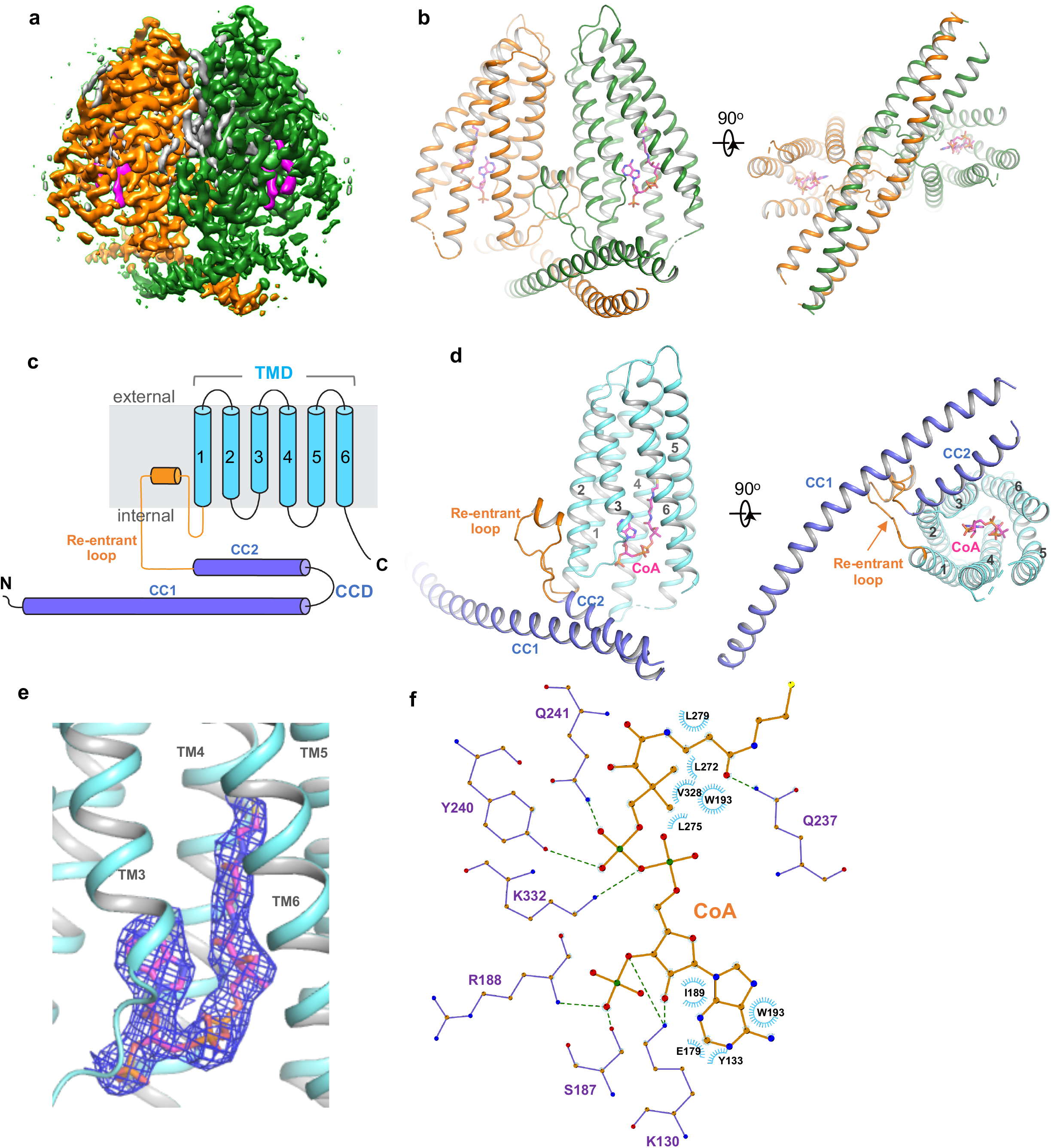
Overall structure of TMEM120A. **(a)** Side view of 3-D reconstruction of TMEM120A. Channel subunits are colored individually with bound substrate density in purple and lipid density in grey. **(b)** Side and bottom views of cartoon representation of TMEM120A structure. Coenzyme A (CoA) molecules are rendered as sticks. **(c)** Topology and domain arrangement in a single TMEM120A subunit. **(d)** Side and bottom views of a single subunit in a similar orientation as the green-colored subunit in b. **(e)** Zoomed-in view of CoA binding site with its density (blue mesh). **(f)** Schematic diagram detailing the interactions between TMEM120A residues and CoA. Toothed wheels mark the hydrophobic contacts between protein residues and CoA. Dotted lines mark the salt bridges and hydrogen bonds.

Each TMEM120A subunit can be divided into two domains: the N-terminal coiled coil domain (CCD) containing CC1 and CC2 helices, and the C-terminal transmembrane domain (TMD) containing six membrane-spanning helices that form an α-helical barrel (Figure 1c&d). The two domains are connected by a well-structured, membrane penetrating re-entrant loop with a short helix (named re-entrant Helix) on its tip. As the cellular localization of TMEM120A as well as its orientation in the membrane is not clearly defined, we therefore arbitrarily consider the coiled coli side of the protein as the internal side and its opposite as external side in our structural description.

TMEM120A forms a tightly packed dimer with extensive dimerization interactions involving multiple parts of the protein (Figure 2a). Dimerization starts at CCD where the exceptionally long (~60 residues) CC1 helix formed an anti-parallel coiled coil with CC1 from the neighboring subunit. CC2 helix has a length of about 1/3 of CC1 and runs anti-parallel to the C-terminal part of CC1, forming a 3-helix bundle with the coiled coil (Figure 2b). The re-entrant loop of each subunit is tightly wedged between the two TMDs of the dimer at the internal leaflet of the membrane. It mediates another set of extensive dimerization interactions through predominantly van de wall contacts with the re-entrant loop from the neighboring subunit as well as the internal halves of TMDs from both subunits (Figure 2c). This insertion of the re-entrant loops between the two subunits splits apart the two TMDs at the internal leaflet of the membrane and consequently the two TMDs make direct contact only at the external leaflet of the membrane through some hydrophobic residues at the external parts of TM2, TM3 and TM6, rendering the TMEM120A dimer with an arrowhead-shaped transmembrane region (Figure 2d). Thus, the extensive dimerization of TMEM120A involving virtually every part of the protein implies that the protein has to function as a dimer.

**Figure 2.**
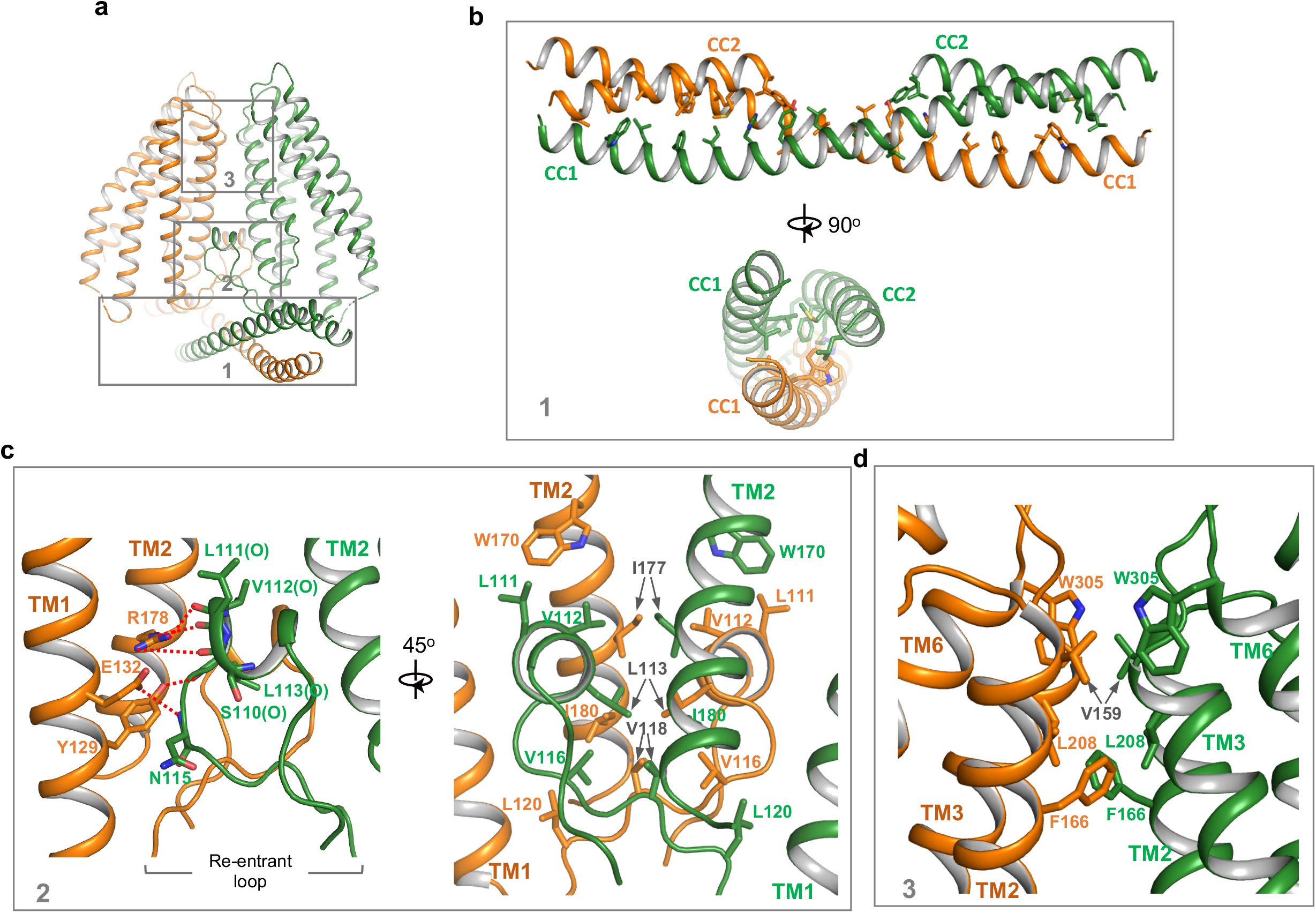
Dimerization of TMEM120A. **(a)** Extensive dimerization interactions occur in three boxed regions: CCD (box 1), re-entrant loop (box 2), and external side of TMD (box 3). **(b)** Zoomed-in view of dimerization interactions at CCD. Residues that participate in the inter-subunit contact are: W16, L19, F23, 126, H30, Y33, L37, L40, L43, I51, L58 and L61 in CC1; L83, M87, L93, F94, M97, Y100 and L101 in CC2. **(c)** Zoomed-in view of dimerization at the re-entrant loop. Shown in left panel are the inter-subunit hydrogen bonding interactions between R178 side chain and the carbonyl oxygen atoms of S110, L111 and V112; between Y129 side chain and the carbonyl oxygen of L113; between the side chains of E132 and N115. Shown in right panel are the intersubunit hydrophobic contacts between the two re-entrant loops and between the re-entrant loop and TMs1-2 of neighboring subunit. **(d)** Zoomed-in view of the inter-subunit hydrophobic contacts between the two TMDs.

### 2. Structural similarities between TMEM120A and ELOVL fatty acid elongase

The overall structure TMEM120A shows no clear feature of a channel protein and has no discernible ion conduction pathway. We performed DALI search and identified the human ELOVL7 structure to share the same fold as TMEM120A at the TMD region (Holm and Rosenstrom, 2010). ELOVL7 is an ER membrane enzyme and belongs to ELOVL family elongases that catalyze the condensation reaction step in the elongation process of very long-chain fatty acid (Jakobsson et al., 2006). ELOVL7 contains seven transmembrane helices and six of them (TMs 2-7) form a 6-TM α-helical barrel that encloses a cytosol-facing pocket where a condensation reaction product of 3-keto acyl-CoA is bound (Nie et al., 2020) (Figure 3a). Despite low sequence homology, the 6-TM barrel structure of ELOVL7 is strikingly similar to that of TMEM120A (TMs 1-6) with a main chain RMSD of about 2.5 Å between their barrel-forming 6-TM helices (Figure 3b). Remote homology at the TMD region between TMEM120A and ELOVL family elongases was also detected by the HHpred server for remote protein homology detection and structure prediction (Soding et al., 2005).

**Figure 3.**
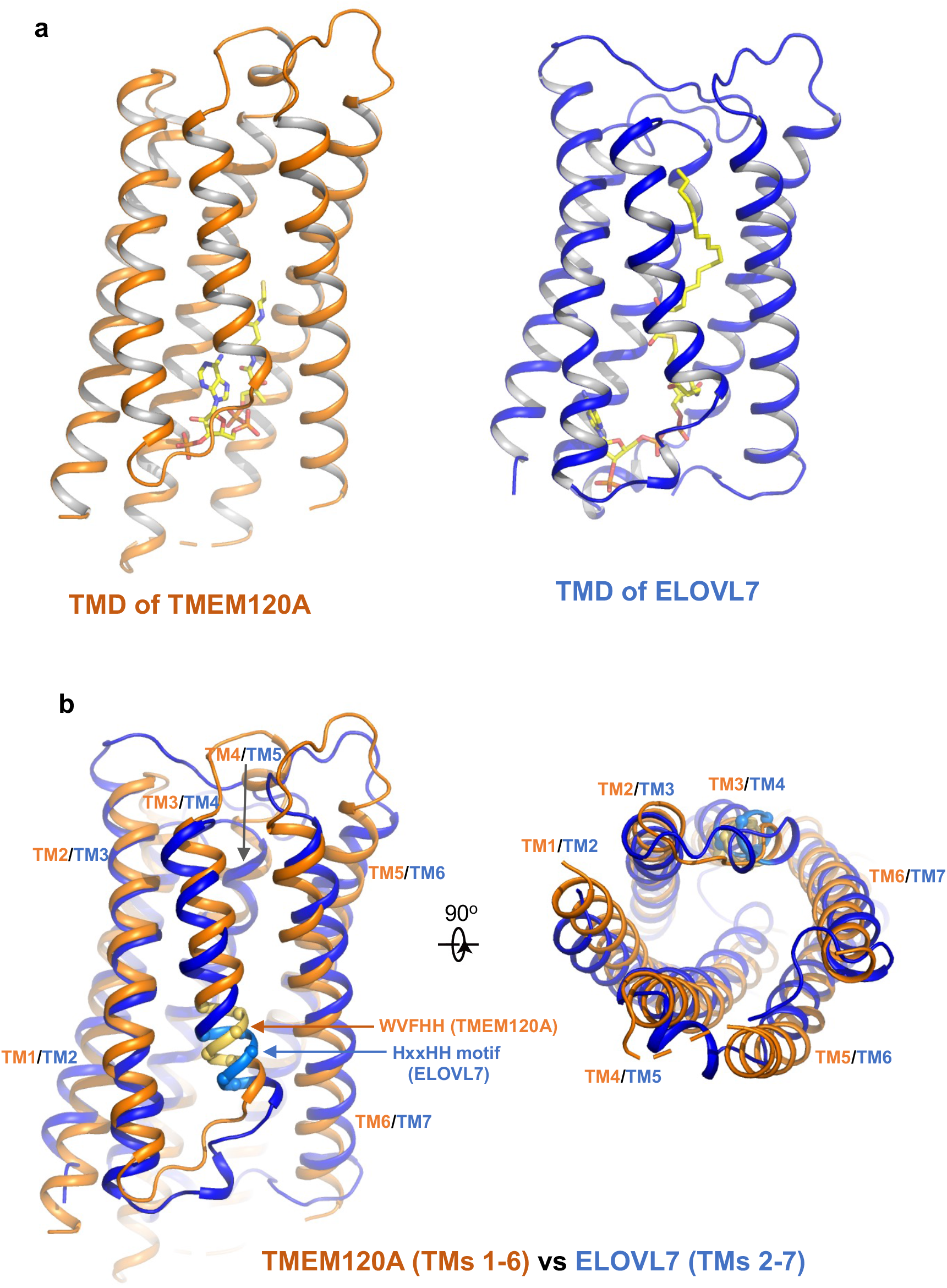
Structural comparison between TMEM120A and ELOVL7 elongase. **(a)** Structures of the 6-TM α-barrel TMDs from TMEM120A (TMs 1-6, left) and ELOVL7 elongase (TMs 2-7, right). CoA in TMEM120A and 3-keto acyl-CoA in ELOVL7 are rendered as sticks. **(b)** Structural comparison between the 6-TM barrels from TMEM120A (orange) and ELOVL7 (blue) in side view (left) and bottom view (right). HxxHH motif in ELOVL7 is colored in cyan. The WVFHH sequence of TMEM120A at the equivalent location is colored in yellow.

ELOVL elongases contain a highly conserved multi-histidine motif (HxxHH) important for their catalytic activity. In ELOVL7 structure, this motif is located at the beginning of TM4 with a sequence of HVFHH (Nie et al., 2020) (Figure 3b). Interestingly, TMEM120A has a sequence of WVFHH at the equivalent location of the 6-TM barrel (the beginning of TM3), almost identical to the histidine motif of ELOVL7. Furthermore, ELOVL elongases bind CoA derivatives as substrates or products and in the ELOVL7 structure a bound 3-keto acyl-CoA product is identified in the deep pocket of the 6-TM barrel (Nie et al., 2020). In TMEM120A structure, we also observe a piece of well-resolved electron density in the pocket of the 6-TM barrel that fits well with a CoA molecule (Figure 1e). Indeed, this bound ligand was confirmed to be CoA by other biochemical assays as discussed in the following section.

### 3. CoA binding in TMEM120A

To confirm the presence of CoA in the purified protein, we measured the CoA level in the protein sample using a commercially available CoA assay kit (MAK034, Sigma-Aldrich). In this assay, CoA in the sample is used to develop products that can react with OxiRed Probe and generate color (OD at 570 nm) and fluorescence (Ex=535nm/Em=587nm). The colorimetric or fluorometric measurement is then used to quantify CoA in the sample by comparing to a CoA standard curve. We used the fluorometric measurement to quantify the CoA level in our sample. As shown in Figure 4a, an assay of 4 mg/ml purified TMEM120A protein sample (~ 0.1 mM, calculated based on OD280) at various volumes yielded a CoA concentration of about 0.15 mM in the sample, matching reasonably well to the calculated concentration of CoA with 1:1 protein/ligand ratio (Figure 4a).

**Figure 4.**
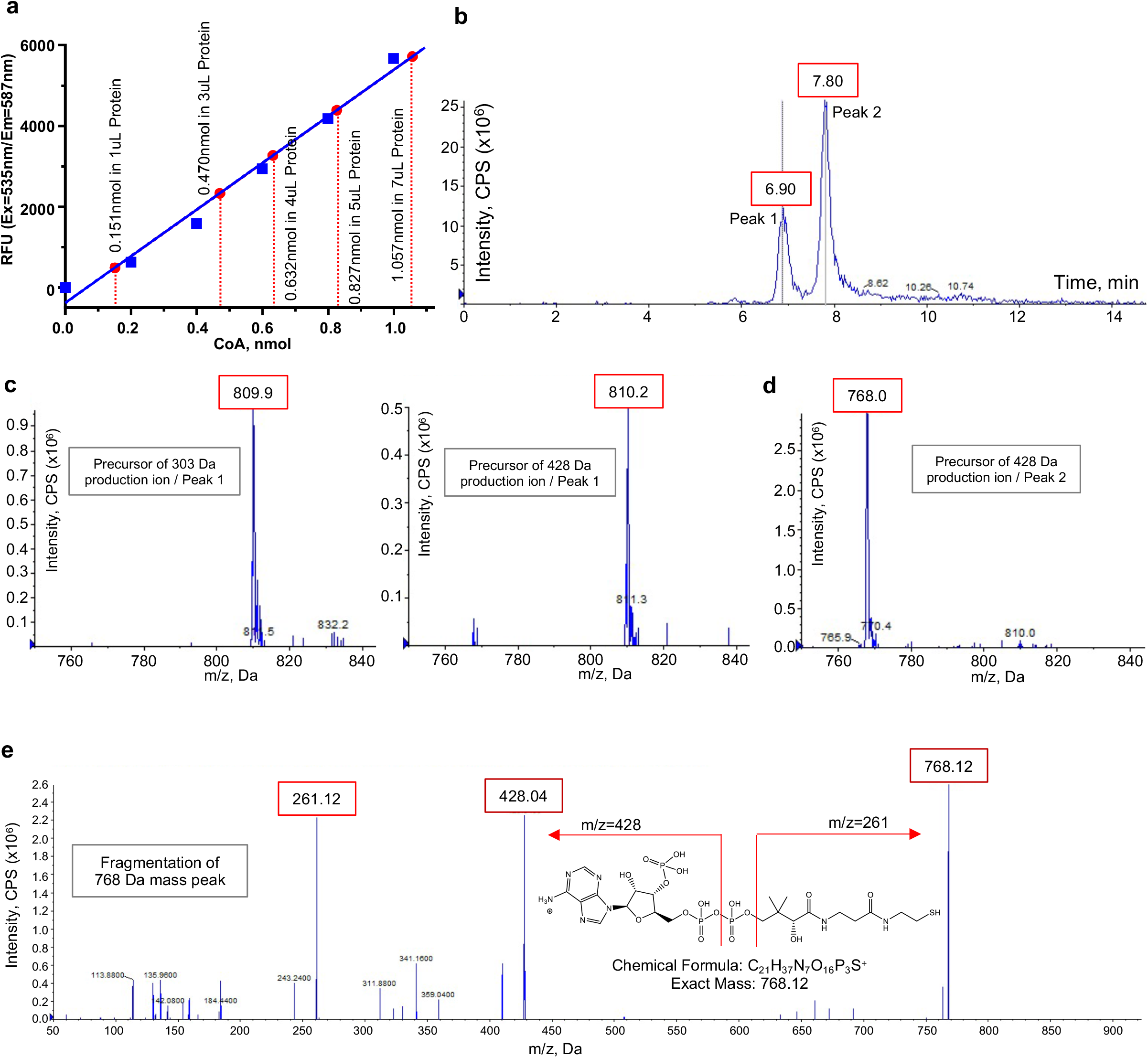
Biochemical and mass spectrometry assay of CoA in TMEM120A. **(a)** CoA assay. Blue standard curve is obtained from fluorometric measurements of various amount of pure CoA provided in the assay kit. Red dots mark the measured CoA contents in 1, 3, 4, 5, and 7 μL of protein samples. The measure CoA concentration in 4mg/mL protein sample is 0.1564±0.0027 mM (mean±SEM, n=5). **(b)** LC separation of extracted substrates from TMEM120A protein sample in LC-MS/MS. **(c)** Precursor ion scan of peak 1 effluent using 303Da (left) and 428Da (right) fragments. **(d)** Precursor ion scan of peak 2 effluent using 428Da fragments. **(e)** Fragmentation (product ion scan) of the 768 Da mass peak.

We also identified the bound CoA in TMEM120A sample using mass spectrometry analysis. In this experiment, the bound substrate was extracted by precipitating the purified protein in high methanol and the substrate mass was analyzed using liquid chromatography with tandem mass spectrometry (LC-MS/MS). Two major peaks with retention time of about 6.9 min and 7.8 min were observed in LC separation (Figure 4b). The first peak at 6.9 min was consistent with the elution volume of acetyl-CoA. In mass spectrometry data acquisition, we performed precursor ion scan for peak 1 effluent using the two main product ions of acetyl-CoA fragmentation (303Da and 428 Da) and identified the precursor ion with the same mass as acetyl-CoA (m/z=810 Da) for both fragments, suggesting the presence of acetyl-CoA in our protein sample (Figure 4c). Peak 2 has a much higher intensity and its effluent has the mass of CoA (m/z=768 Da) in the precursor ion scan using the fragment of 428 Da (a main product ion of CoA) (Figure 4d). The product ion scan of the 768 Da mass peak yielded the same fragmentation pattern as CoA, confirming the identity of CoA in peak 2 (Figure 4e). Thus, mass spectrometry analysis identified both CoA and acetyl-CoA in our protein sample. Combining our structural observation with the biochemical CoA assay and mass spectrometry analysis, we suspect that CoA is likely the main substrate in the purified protein sample.

## Summary

Here we present some structural and biochemical analyses of membrane protein TMEM120A which forms a tightly packed dimer. The transmembrane domain of each TMEM120A subunit forms a 6-TM helical barrel where a CoA molecule can bind. While TMEM120A was recently proposed to function as a mechanosensitive channel, its structure shows no clear feature of an ion channel. Despite low sequence homology, TMEM120A structure shares some striking similarities to ELOVL7, an ER membrane elongase for very long-chain fatty acid. Firstly, TMDs of both proteins contain a 6-TM α-barrel with very similar topology and architecture. Secondly, both proteins can bind CoA or CoA derivative in the pocket of the 6-TM barrel. Thirdly, the conserved HxxHH motif important for the catalytic activity of ELOVL elongase is also present at the equivalent location in TMEM120A. Although the exact physiological function of TMEM120A remains to be determined, its similarity to ELOVL fatty acid elongase is unlikely to be coincidental and may imply enzymatic function of TEME120A for fat metabolism.

## Materials and Methods

### Key resources table

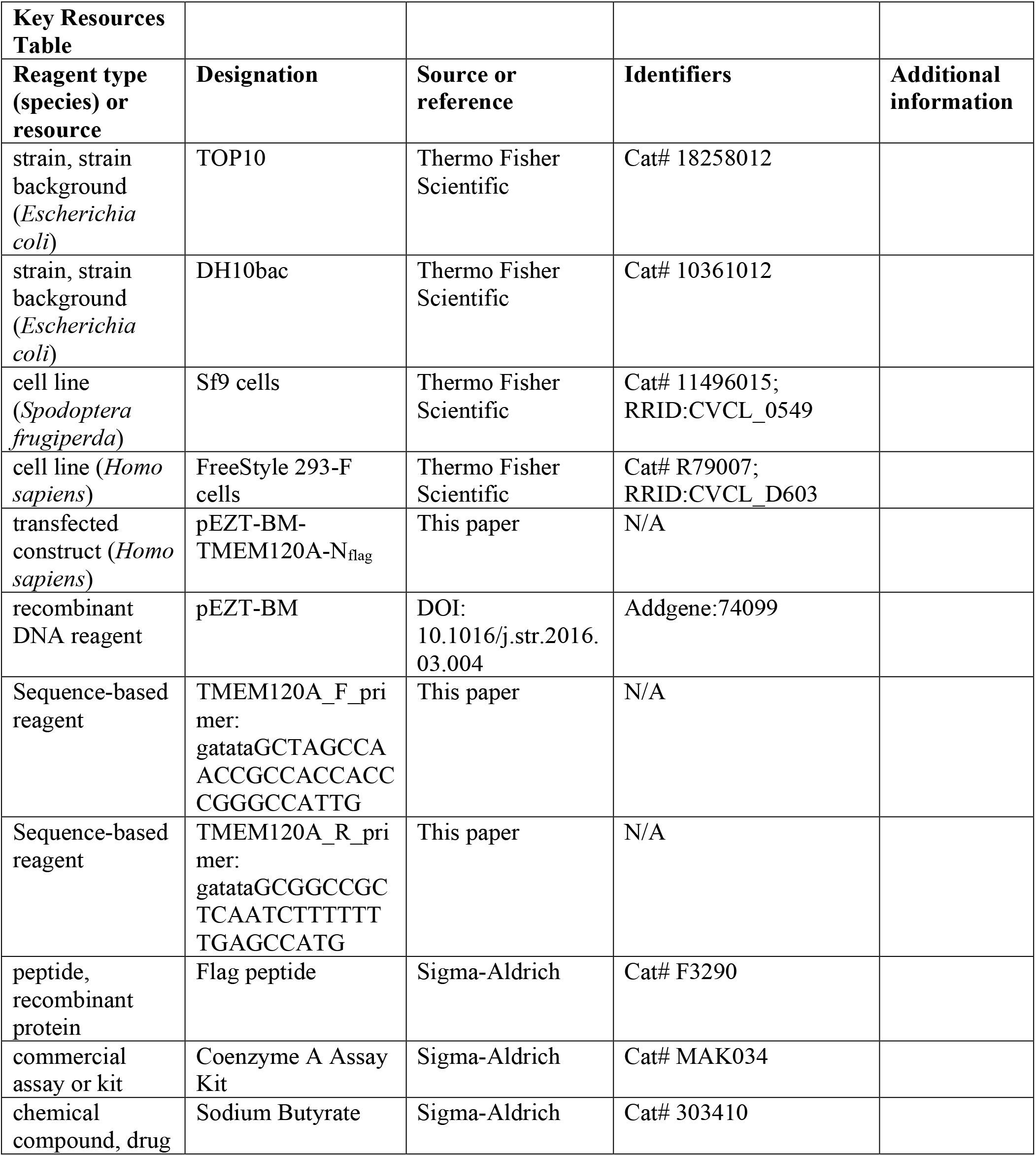
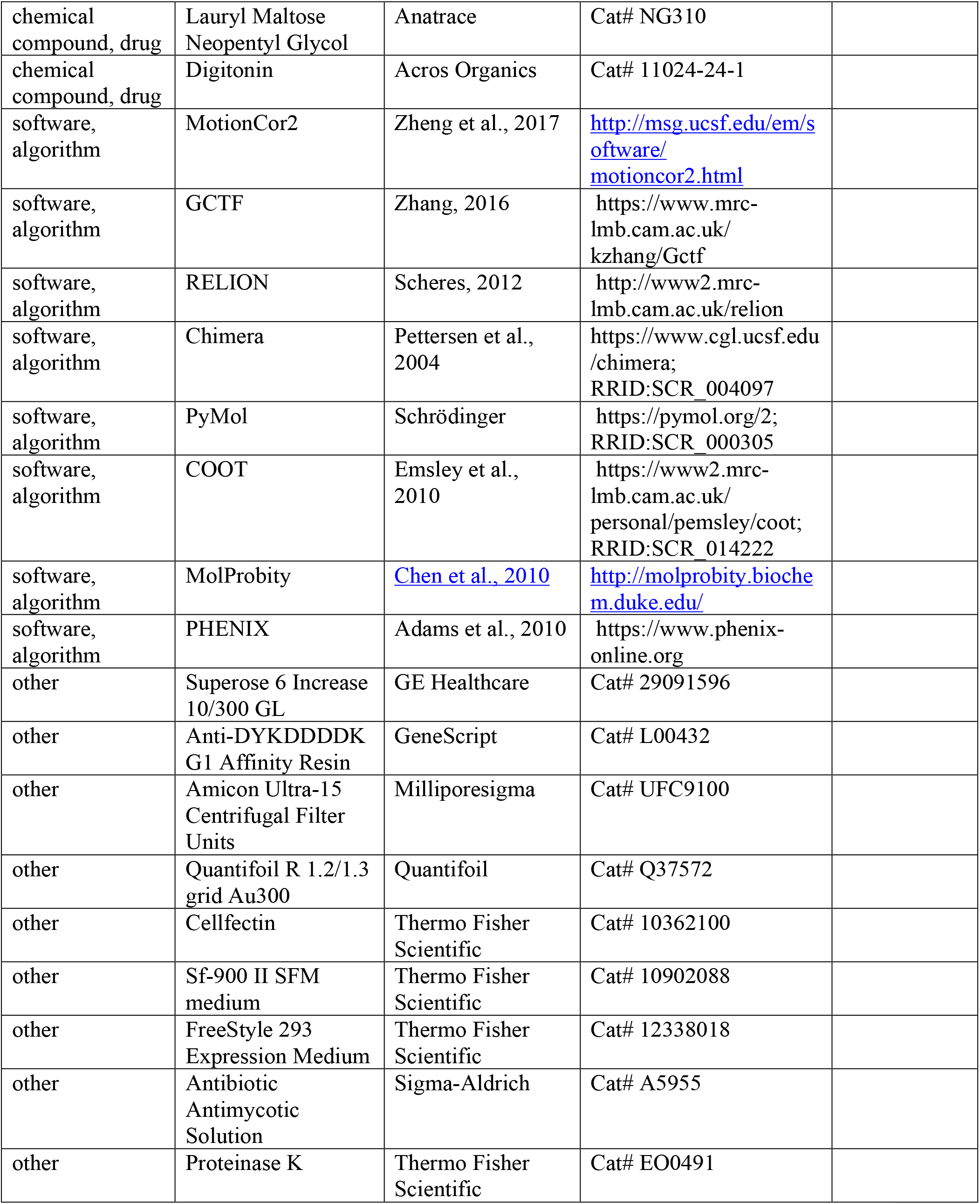

### Protein expression and purification

Full-length Homo sapiens TMEM120A (HsTMEM120A, NCBI accession: NP_114131.1) was cloned into a modified pEZT-BM vector with an N-terminal Flag tag (Morales-Perez et al., 2016) and heterologously expressed in HEK293F cells using the BacMam system (Thermo Fisher Scientific). Bacmids were synthesized using E. coli DH10bac cells (Thermo Fisher Scientific) and baculoviruses were produced in Sf9 cells using Cellfectin II reagent (Thermo Fisher Scientific). For protein expression, cultured HEK293F cells were infected with the baculoviruses at a ratio of 1:40 (virus:HEK293F, v/v) and supplemented with 10mM sodium butyrate to boost protein expression level. Cells were cultured in suspension at 37°C for 48 hr and then harvested by centrifugation at 4,000 g for 15 min. All purification procedures were carried out at 4°C unless specified otherwise. The cell pellet was resuspended in buffer A (35 mM HEPES pH 7.4, 300 mM NaCl) supplemented with protease inhibitors (2 μg/ml DNase, 0.5 μg/ml pepstatin, 2 μg/ml leupeptin, and 1 μg/ml aprotinin and 0.1 mM PMSF). After homogenization by sonication, HsTMEM120A was extracted with 1% (w/v) Lauryl Maltose Neopentyl Glycol (LMNG, Anatrace) by gentle agitation for 2 hr. After extraction, the supernatant was collected by centrifugation at 40,000 g for 30 min and incubated with anti-Flag G1 affinity resin (Genescript) by gentle agitation for 1 hr. The resin was then collected on a disposable gravity column (Bio-Rad) and washed with 20 column volume of Buffer A supplemented with 0.05% (w/v) LMNG followed by 20 column volume of Buffer B (25 mM HEPES pH 7.4, 150 mM NaCl) supplemented with 0.06% (w/v) Digitonin (ACROS Organics). TMEM120A was eluted in Buffer B with 0.06% (w/v) Digitonin and 0.2 mg/ml Flag peptide. The protein eluate was concentrated and further purified by size-exclusion chromatography on a Superdex200 10/300 GL column (GE Healthcare) in Buffer B with 0.06% (w/v) Digitonin. The peak fractions were collected and concentrated to 5 mg/ml for cryo-EM analysis.

HEK293F cells (RRID: CVCL_D603) were purchased from and authenticated by Thermo Fisher Scientific. The cell lines were tested negative for mycoplasma contamination.

### Cryo-EM data acquisition

Purified HsTMEM120A at 5 mg/ml was applied to a glow-discharged Quantifoil R1.2/1.3 300-mesh gold holey carbon grid (Quantifoil, Micro Tools GmbH, Germany), blotted under 100% humidity at 4°C and plunged into liquid ethane using a Mark IV Vitrobot (FEI).

Cryo-EM data were acquired on a Titan Krios microscope (FEI) at the HHMI Janelia Cryo-EM Facility operated at 300 kV with a K3 Summit direct electron detector (Gatan), using a slit width of 20 eV on a GIF-Quantum energy filter. Images were recorded with Serial EM in superresolution counting mode with a super resolution pixel size of 0.422 Å. The defocus range was set from −0.9 to −2.2 μm. Each movie was dose-fractionated to 60 frames under a dose rate of 9.2 e-/pixel/s using CDS (Correlated Double Sampling) mode of the K3 camera, with a total exposure time of 4.646 s, resulting in a total dose of 60 e-/Å2.

### Cryo-EM Image processing

Movie frames were motion corrected and binned two times and dose-weighted using MotionCor2 (Zheng et al., 2017). The CTF parameters of the micrographs were estimated using the GCTF program (Zhang, 2016). The rest of the image processing steps was carried out using RELION 3.1 (Nakane et al., 2020; Scheres, 2012; Zivanov et al., 2018). The map resolution was reported according to the gold-standard Fourier shell correlation (FSC) using the 0.143 criterion (Henderson et al., 2012). Local resolution was estimated using Relion.

Aligned micrographs were manually inspected to remove those with ice contamination and bad defocus. Particles were selected using Gautomatch (Kai Zhang, http://www.mrc-lmb.cam.ac.uk/kzhang/) and extracted using a binning factor of 3 (box size was 96 pixels after binning). 2D classification was performed in Relion 3.1. Selected particles after 2D classification were subjected to one around of 3D classification. Ab initio model was generated in Relion 3.1 and used as the reference for this 3D classification. Classes that showed similar structure features were combined and subjected to 3D auto-refinement and another round of 3D classification without performing particle alignment using a soft mask around the protein portion of the density. The best resolving classes were re-extracted with the original pixel size and further refined. Beam tilt, anisotropic magnification, and per-particle CTF estimations and Bayesian polishing were performed in Relion 3.1 to improve the resolution of the final reconstruction.

### Model building, refinement and validation

EM map of HsTMEM120A is of high quality for de novo model building in Coot (Emsley et al., 2010). The model was manually adjusted in Coot and refined against the map by using the real space refinement module with secondary structure and non-crystallographic symmetry restraints in the Phenix package (Adams et al., 2010).

The final structural model of each subunit contains residues 8-69, 80-255 and 261-335. Residues of 1-7, 70-79, 256-260 and 335-343 were disordered in the structure. The statistics of the geometries of the models were generated using MolProbity (Chen et al., 2010). All the figures were prepared in PyMol (Schrödinger, LLC.) and UCSF Chimera (Pettersen et al., 2004).

The multiple sequence alignments were performed using the program Clustal Omega (Sievers et al., 2011).

### Coenzyme A Quantification Assay

HsTMEM120A was purified using the same protocol as described above. To release any bound CoA substrate from the protein, HsTMEM120A was subjected to protease digestion with 1mg/ml proteinase K at 37°C for 1h (Thermo Scientific; EO0491). 0.2% SDS was added to the digestion solution to stimulate the activity of proteinase K. After digestion, proteinase K was denatured by incubating the sample at 70°C for 7min.

CoA levels in the protein solution after proteinase K digestion were quantified using a commercial CoA assay kit according to manufacturer’s protocol (Sigma-Aldrich; MAK034). CoA concentration is determined by an enzymatic assay, in which a colored product is developed and the colorimetric (OD at 570 nm) or fluorometric (Ex=535nm/Em=587 nm) measurement of the product is proportional to the amount of CoA in the sample. We used fluorometric measurement in our assay for CoA quantification and its concentration was determined by comparing to a standard curve plotted from using the pure CoA standard in the assay.

### Liquid chromatography-mass spectrometry (LC-MS/MS) analysis

HsTMEM120A sample for mass spectrometry (MS) assay was purified using the similar protocol as described above with slight modification. The collected anti-Flag G1 affinity resin was washed with 20 column volume of Buffer C (25 mM HEPES pH 7.4, 180 mM NaCl) supplemented with 0.01% (w/v) LMNG. HsTMEM120A was eluted in Buffer C with 0.01% (w/v) LMNG and 0.2 mg/ml Flag peptide. The protein eluate was concentrated and further purified by size-exclusion chromatography on a Superdex200 10/300 GL column (GE Healthcare) in Buffer C with 0.005% (w/v) LMNG. The peak fractions were collected and concentrated to 13 mg/ml for MS analysis.

To extract the bound CoA substrate, the protein was precipitated by adding 640uL of MeOH (LC-MS grade) to 160uL of concentrated TMEM120A sample (13 mg/ml) followed by 30 sec of vortex. The sample was kept in −20 °C freezer for 20 min before collecting the supernatant by centrifugation (16,400g) for 10 min at 4 °C. The supernatant was filtered (0.2 micron PVDF filter) before MS analysis.

LC-MS/MS analysis was conducted using a SCIEX QTRAP 6500+ mass spectrometer coupled to a Shimadzu HPLC (Nexera X2 LC-30AD). The ESI source was used in positive ion mode. The ion spray needle voltage was set at 5500 V. HILIC chromatography was performed using a SeQuant® ZIC-pHILIC 5μm polymer 150 x 2.1 mm PEEK coated HPLC column (Millipore Sigma, USA). The column temperature, sample injection volume, and flow rate were set to 45°C, 5 μL, and 0.15 mL/min, respectively. HPLC solvent and gradient conditions were as follows: Solvent A: 20 mM ammonium carbonate, 0.1% ammonium hydroxide. Solvent B: 100% acetonitrile. Gradient conditions were 0 min: 20% A + 80% B, 20 min: 80% A + 20% B, 22 min 20% A + 80% B, 34 min: 20% A + 80% B. Total run time: 34 mins. Flow was diverted to waste for the first 5 min and after 16 min.

A precursor ion (PI) scan in the range of 750-1250 Da was used to identify parent ions that yielded two product ions of 303 Da and 428 Da which are characteristic of acetyl-CoA. This strategy was applied to monitor the presence of acetyl-CoA and other acyl-CoAs in the sample. An acetyl-CoA standard was used to confirm retention time and fragmentation to product ions. In addition, a EMS-IDA-EPI scan was used to fragment the mass peak observed at 768 Da, which was subsequently assigned as CoA. Data were analyzed using Analyst 1.7.1 software.

## Acknowledgements

Single particle cryo-EM data were collected at the University of Texas Southwestern Medical Center Cryo-EM Facility that is funded by the CPRIT Core Facility Support Award RP170644 and the Howard Hughes Medical Institute Janelia Cryo-EM Facility. We thank Rui Yan at the Janelia Cryo-EM Facility for help in microscope operation and data collection. This work was supported in part by the Howard Hughes Medical Institute (Y.J. and N.G.) and by grants from the National Institute of Health (GM079179 to Y. J.), the Welch Foundation (Grant I-1578 to Y. J.).

## Authors Contributions

J.X. prepared the samples; J.X. and Y.H. performed EM data acquisition, image processing and structure determination; H.B. and B.P.T performed mass spectrometry analysis; J.P. and N.G. performed computational analysis of TMEM120 family proteins; J.W. performed molecular dynamic simulations; W.Z. performed electrophysiology recordings; J.X., Y.J., H.B. and B.P.T participated in research design, data analysis, discussion and manuscript preparation.

## Contact for Reagent and Resource Sharing

The cryo-EM density map and the atomic coordinates of the human TMEM120A have been deposited in the Electron Microscopy Data Bank under accession numbers EMD**-**24230 and the Protein Data Bank under accession numbers 7N7P, respectively. Further information and requests for resources and reagents should be directed to and will be fulfilled by Lead Contact, Youxing Jiang

---

**Figure 1—figure supplement 1.**
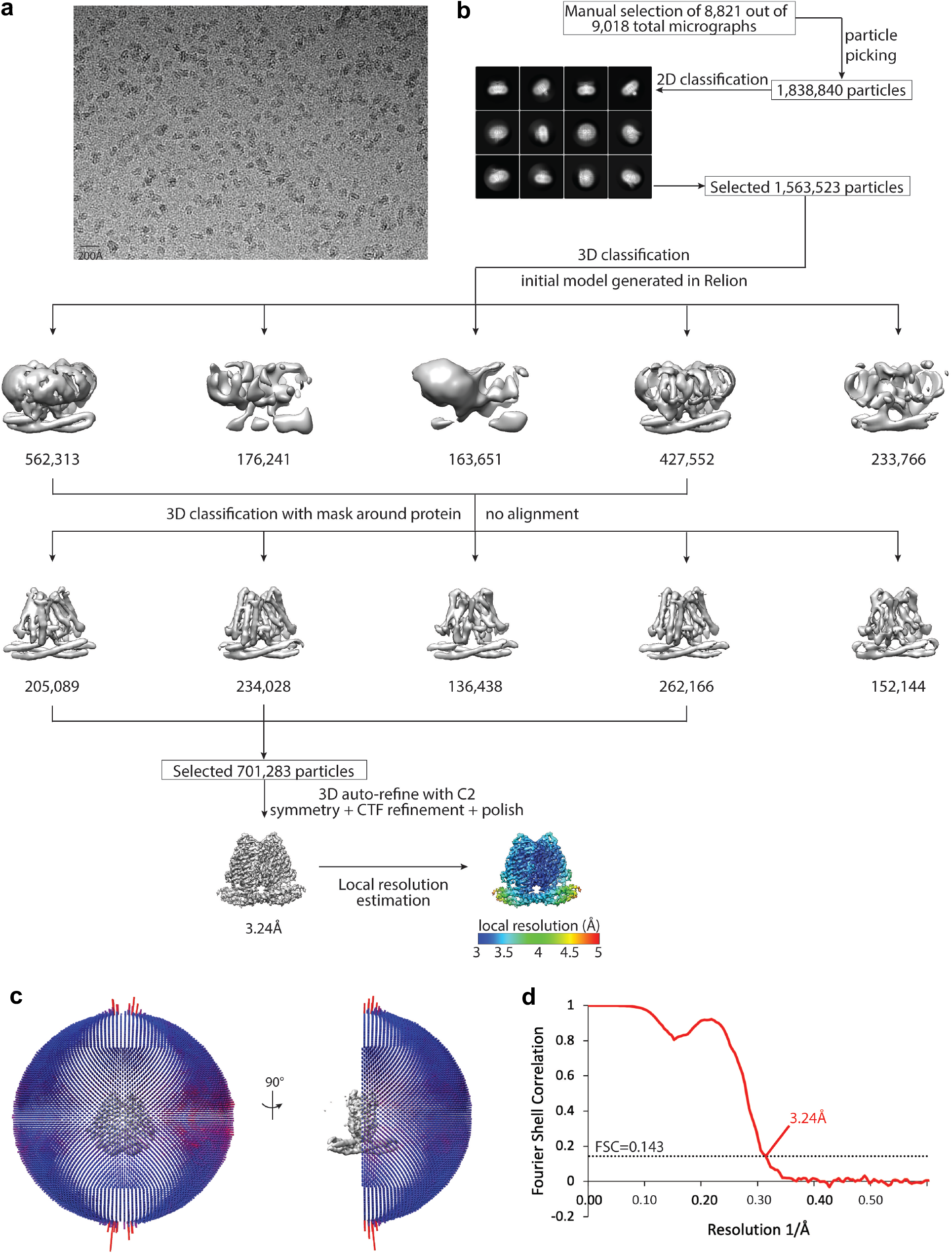
Cryo-EM data processing scheme of the cGMP-bound human TMEM120A. **(a)** A representative micrograph. Scale bar is at 20 nm. **(b)** Flow chart of the cryo-EM data processing procedure. Selected 2D class averages are shown. The particle numbers are indicated under the corresponding 3D classes. **(c)** Euler angle distribution of particles used in the final three-dimensional reconstruction. **(d)** Fourier Shell Correlation curves showing the overall resolution of 3.24 Å at FSC=0.143.

**Figure 1—figure supplement 2.**
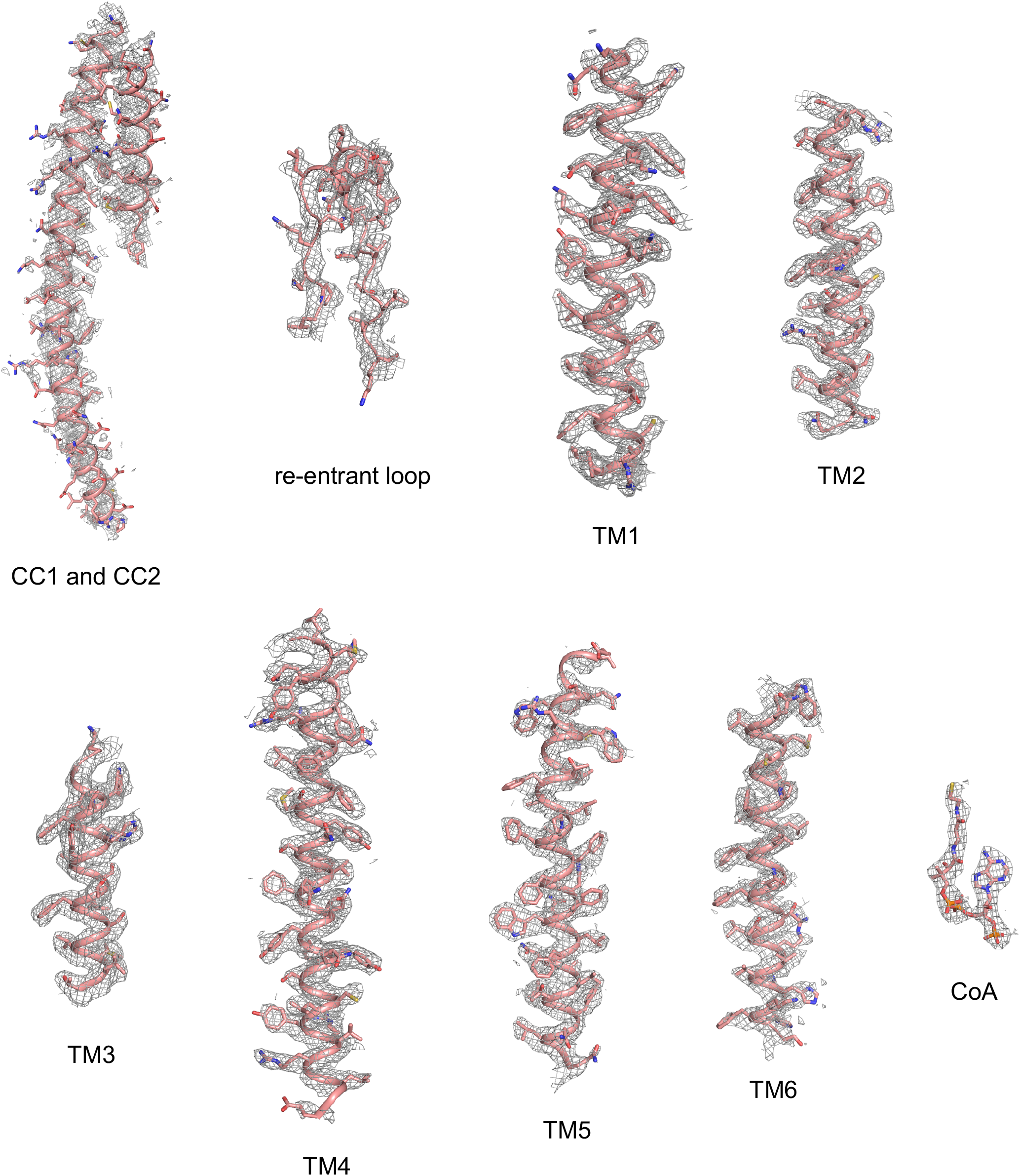
Sample density maps of human TMEM120A at various regions. Maps of CC1 and CC2 are contoured at 4.5 σ. All other maps are contoured at 6.0 σ.

**Figure 1—figure supplement 3.**
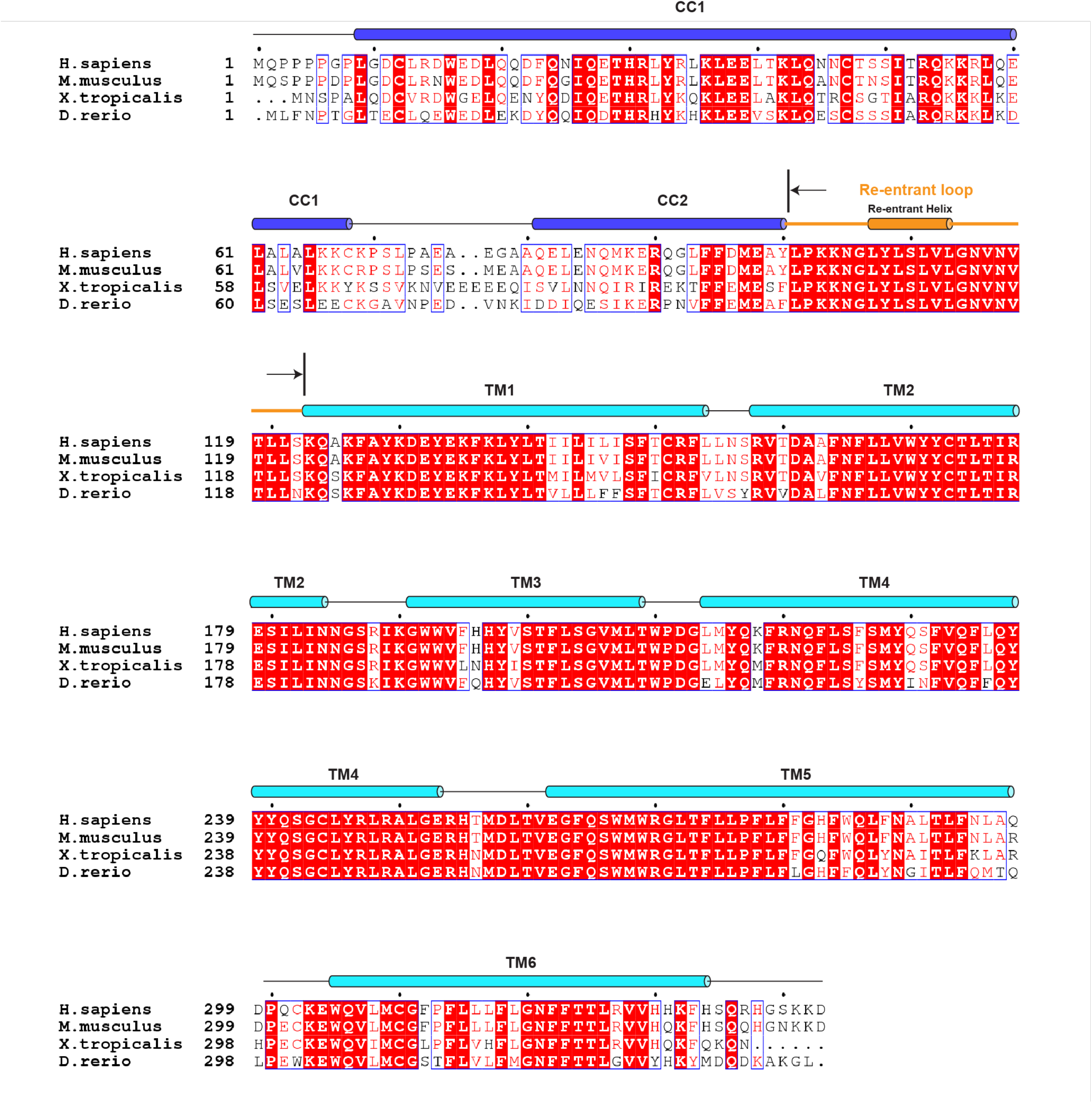
Sequence alignment of vertebrate TMEM120A. Secondary structure assignments are based on the structure of human TMEM120A.

**Figure 1—source data 1.**
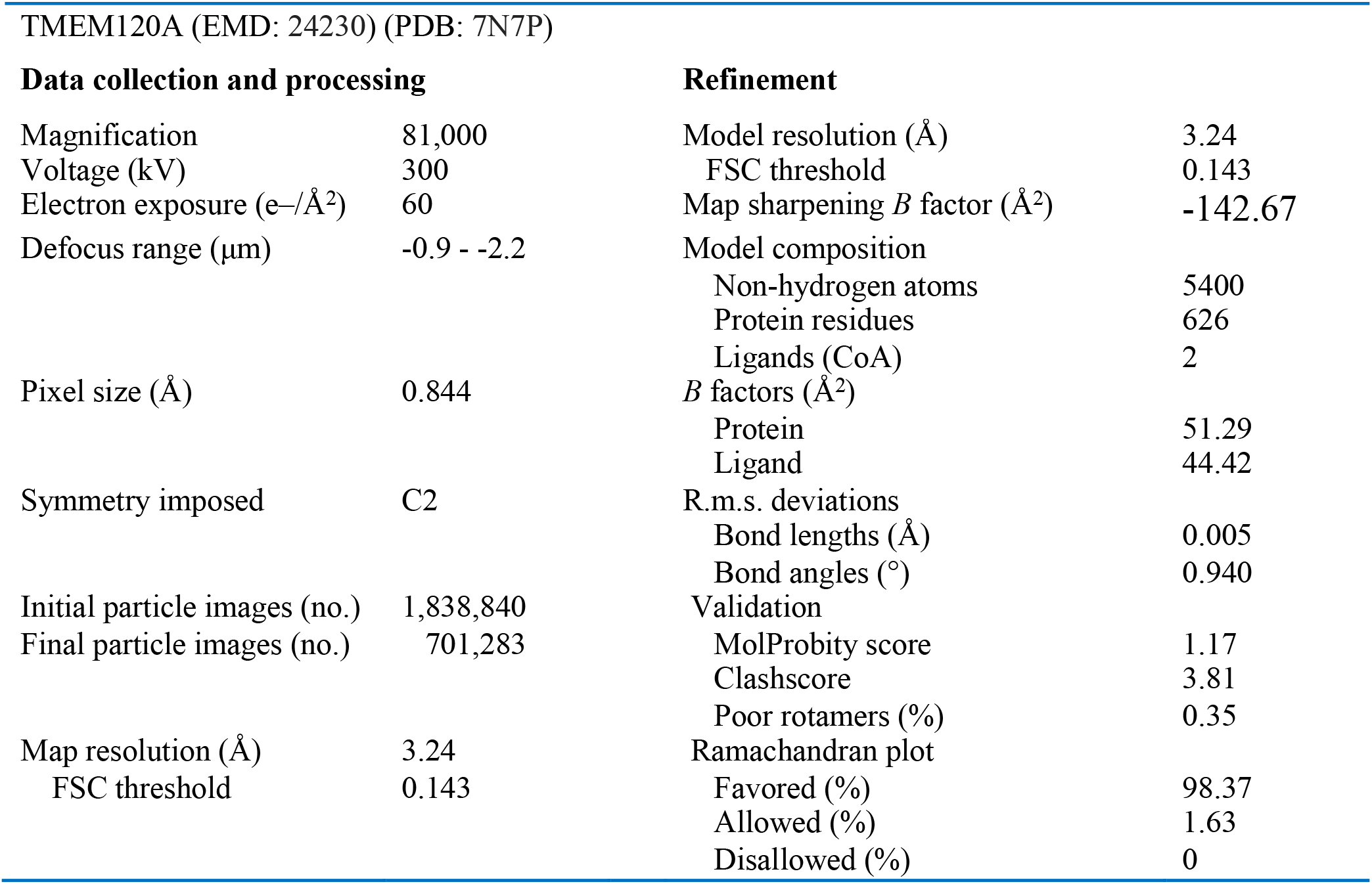
Cryo-EM data collection and model statistics.

